# AAV-mediated gene augmentation therapy restores critical functions in mutant iPSC-derived PRPF31^+/-^ cells

**DOI:** 10.1101/729160

**Authors:** Elizabeth M. Brydon, Revital Bronstein, Adriana Buskin, Majlinda Lako, Eric A. Pierce, Rosario Fernandez-Godino

## Abstract

Retinitis pigmentosa (RP) is the most common form of inherited vision loss and is characterized by degeneration of retinal photoreceptor cells and the retinal pigment epithelium (RPE). Mutations in pre-mRNA processing factor 31 (*PRPF31*) cause dominant RP via haploinsufficiency with incomplete penetrance. There is good evidence that the diverse severity of this disease is a result of differing levels of expression of the wild type allele among patients. Thus, we hypothesize that *PRPF31*-related RP will be amenable to treatment by adeno-associated virus (AAV)-mediated gene augmentation therapy. To test this hypothesis, we used induced pluripotent stem cells (iPSC) with mutations in *PRPF31* and differentiated them into RPE cells. The mutant *PRPF31* iPSC-RPE cells recapitulate the cellular phenotype associated with the PRPF31 pathology, including defective cell structure, diminished phagocytic function, defects in ciliogenesis, and compromised barrier function. Treatment of the mutant *PRPF31* iPSC-RPE cells with AAV-*PRPF31* restored normal phagocytosis and cilia formation, and partially restored structure and barrier function. These results provide proof-of concept that AAV-based gene therapy can be used to treat patients with *PRPF31*-related RP.

## INTRODUCTION

With a prevalence rate of approximately 1 in 3500, retinitis pigmentosa (RP) is the most common form of inherited blindness, and can be inherited in autosomal dominant, autosomal recessive, or X-linked patterns ^1^. The general pathology of RP is degeneration of the retinal pigment epithelium (RPE) and photoreceptor cells, sometimes leading to complete vision loss ^2^. The most common causes of autosomal dominant RP are mutations in rhodopsin, followed by mutations in precursor messenger RNA (pre-mRNA) processing factors *PRPF3, PRPF4, PRPF6, PRPF8, PRPF31*, and *SNRNP200* ^3–7^. Among these, mutations in *PRPF31* are the most common, and are estimated to account for approximately 10% of dominant RP ^8^. *PRPF31* encodes a ubiquitously expressed splicing factor, which binds and stabilizes the tri-snRNP U4/U6-U5 ribonucleoprotein complex ^9–11^. It remains unclear why mutations in ubiquitously expressed splicing factors result in disease specific to the retina. Data obtained from studies of *PRPF31*-associated disease suggest that alterations in RNA splicing underlie these forms of inherited retinal degeneration (IRD) ^12^.

RP from mutations in *PRPF31* stems from nonsense mutations, large-scale deletions, and premature stop codons affecting one allele ^10^. These mutations create null alleles and cause disease via haploinsufficiency. Complete loss of PRPF31 function results in embryonic lethality ^10^. Since mutations in *PRPF31* cause disease via haploinsufficiency, it is a dominant disease that is a good candidate for treatment via gene augmentation therapy. Further, evidence from studies of the reduced penetrance of disease observed in some families with *PRPF31*-associated retinal degeneration shows that increased expression of *PRPF31* from the wild-type allele can reduce disease severity ^13–15^. For gene-based therapies, adeno-associated viral (AAV) vectors are at the forefront, since they are known to be non-pathogenic while simultaneously staying successful at penetrating cell membranes and mostly evading the immune system ^16^. Last year, the first FDA-approved gene therapy treatment for inherited retinal diseases was successfully performed in patients with mutations in the retinal pigment epithelium–specific 65-kDa protein (RPE65) gene. Sub-retinal injection of the RPE65-expressing AAV vector restores normal function of this protein and leads to vision improvement ^17^. Stimulated by this initial success, clinical trials of AAV-mediated gene augmentation therapies are in progress for multiple genetic subtypes of inherited retinal degeneration ^18–23^.

Among other functions, the RPE nourishes photoreceptor cells and phagocytoses shed photoreceptor outer segments (POS) ^24^. Mutations in *PRPF31* primarily lead to RPE degeneration in cellular and mouse models of *PRPF31*-linked RP ^9,12,25^. Specifically, RPE cells from *Prpf31* mutant mice show progressive degeneration and a cell-autonomous phagocytic defect associated with decreased binding and internalization of POS that eventually leads to photoreceptor loss ^6^. Since RPE can be derived from induced pluripotent stem cells (iPSC), the RPE pathology associated with mutations in *PRPF31* can be modeled using patient derived iPSC-RPE. Indeed, iPSC-RPE generated from patients with *PRPF31*-associated retinal degeneration show decreased phagocytosis and abnormal cilia growth ^12^. In the studies reported here, we used one of these patient derived iPSC lines and a newly generated *PRPF31*^+/-^ iPSC cell line to test the use of adeno-associated virus (AAV) mediated gene augmentation therapy to treat *PRPF31*-associated retinal degeneration. Data obtained from these studies support the use of AAV-mediated gene augmentation therapy to treat *PRPF31*-associated disease.

## RESULTS

### 1. Generation of iPSC-RPE cells *PRPF31*^+*/-*^ via CRISPR-Cas9 editing

To test AAV-mediated gene augmentation therapy for *PRPF31*-associated IRD, we used iPSC-derived RPE cells from two sources. The first source is iPSC-derived RPE cells from a patient with 11 bp deletion in exon 11 *PRPF31*, causing RP ^12^. RPE cells derived from the same iPSC line in which the 11 bp deletion was corrected were used as controls ^12^. These *PRPF31* mutant iPSC-derived RPE cells reproduce *in vitro* key features associated with *PRPF31* pathology, such as defective splicing, decreased phagocytosis, and shorter cilia ^12^. The second source of iPSCs is wild-type IMR90 iPSC into which we introduced a null allele of *PRPF31* using CRISPR/Cas9 mediated genome editing. To accomplish this modification, we transfected wild type iPSCs with the pSpCas9(BB)2A-EGFP (PX458) plasmid carrying the Cas9 nuclease and a guide RNA (gRNA) targeting exon 7 of PRPF31 (Figure 1). EGFP positive cells were sorted and expanded to generate clonal cell lines. Screening of the clones via PCR and sequencing identified 18/255 clones with mutations in *PRPF31* (8%). The most common indels found in these clones were 4 bp and 10 bp deletions in exon 7 of *PRPF31*, which resulted in frameshift mutations, causing premature stop codons in exon 7 and 8 respectively (Figure 1). Cells transfected with editing reagents, but which did not have a mutation in *PRPF31* were expanded for use as controls. In clones harboring both the 4 bp and 10 bp deletions, the mRNA levels of *PRPF31* were reduced to half compared to counterpart wild type clones (Figure 1B; 2-way ANOVA, p<0.0001).

**Figure 1.**
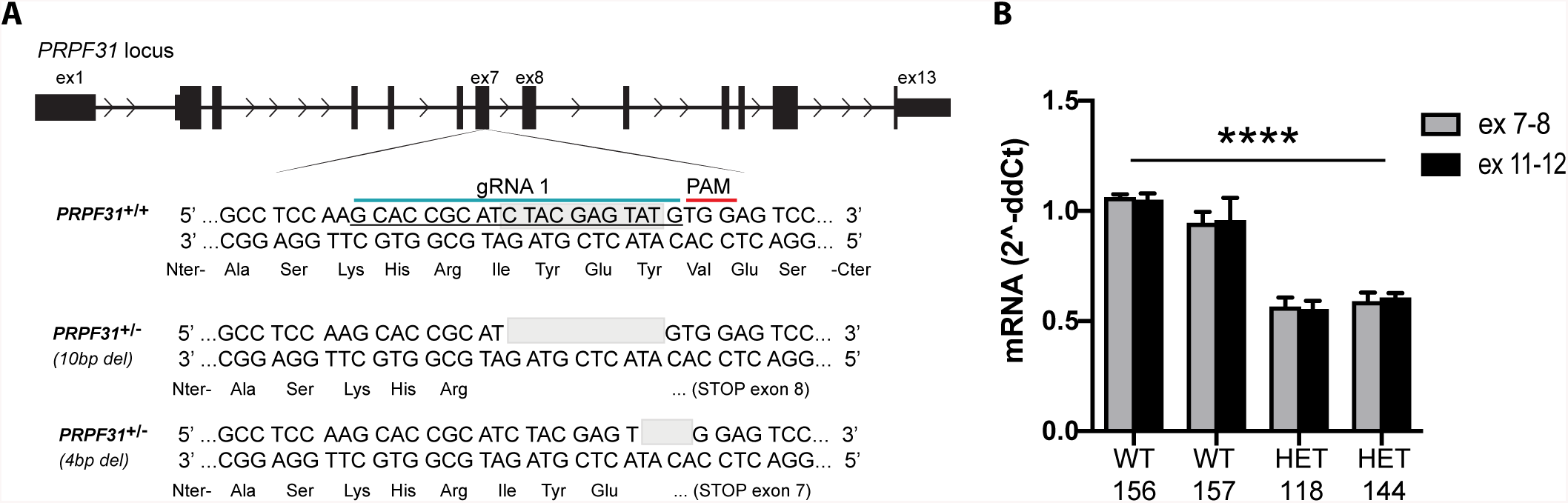
CRISPR-edited iPSC *PRPF31*^+*/-*^. **(A)** Schematic representation of the *PRPF31* locus. 20bp nucleotide gRNA sequence (blue line) followed by PAM (red line) designed to target exon 7. Bottom sequence shows the 10 bp deletion found in clone #144, which was used for differentiation into RPE. **(B)** mRNA levels of *PRPF31* normalized to *GAPDH* measured in triplicate, expressed by CRISPR-edited iPSC *PRPF31*^+*/*+^ (WT) clones 156 and 157, and *PRPF31*^+*/-*^ (HET) mutant clones 118 (4 bp deletion) and 144 (10 bp deletion). The average expression of WT cells was used as value 1 for relative quantification. (2-way ANOVA, ****p<0.0001. Data represented as mean ± SD).

One wild type clone (clone #157) and one clone harboring the 10 bp deletion in one allele of *PRPF31* (clone #144) were chosen for further differentiation into RPE cells, according to a previously established protocol ^26,27^. At passage 2 (p2), iPSC-RPE cells on transwells displayed typical honeycomb morphology, pigmentation, and polarization (Figure 2). The RPE monolayer was formed as shown by the expression of the tight-junction protein ZO-1 (Figure 2B). Successful differentiation into RPE cells was determined through expression of the RPE markers RPE65, TYR (pigmentation enzyme), and RLBP1 (a visual cycle gene), which were not expressed in the iPSCs (Figure 2C). To be functional, the RPE monolayer needs to be highly polarized ^24^. One of the methods to assay RPE polarization is measuring the transepithelial electrical resistance (TER). Despite the normal expression of ZO-1, the engineered iPSC-RPE *PRPF31*^+*/-*^ cells showed significantly lower TER than the counterpart wild type cells (t-test, n=4/genotype. p=0.0009), corroborating results found in patient-derived iPSC-RPE cells ^12^ (Figure 2D).

**Figure 2.**
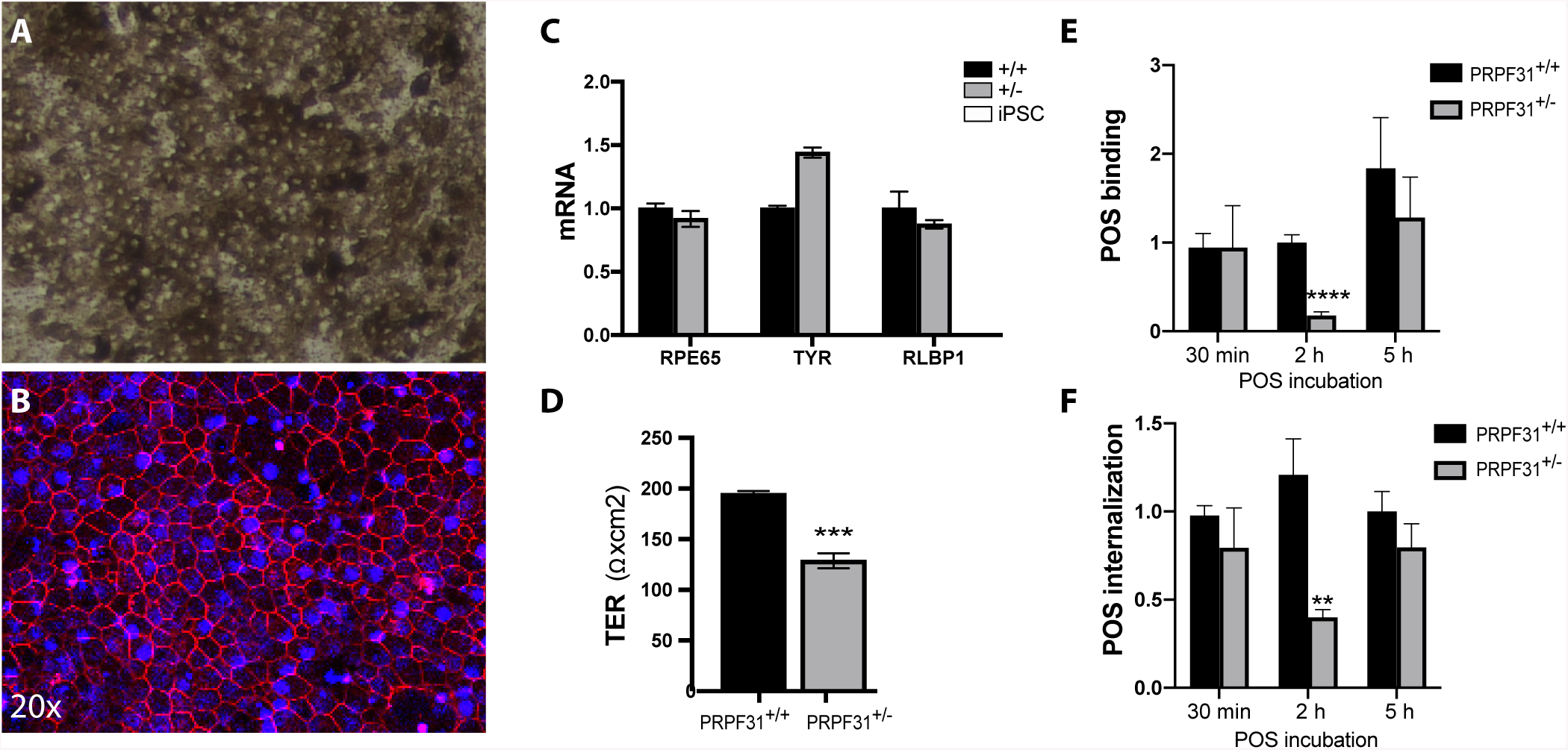
Characterization of the CRISPR-edited iPSC-RPE cell monolayer. **(A)** Brightfield micrograph of mature iPSC-RPE *PRPF31*^+*/-*^ cells on transwells. **(B)** Fluorescent micrograph of mature CRISPR-edited iPSC-RPE *PRPF31*^+*/-*^ cells grown on transwells and immunostained anti-ZO-1 antibody (red) and DAPI (blue). **(C)** RPE markers normalized to *UBE2R2* expressed by mature iPSC-RPE cells wild type *PRPF31*^+*/*+^ (+/+ black bars) and mutant *PRPF31*^+*/-*^ (+/-grey bars) compared to iPSC (white bars, no expression found) (n = 3 replicates/cell type). Data represented as mean ± SD). **(D)** Transepithelial electrical resistance (TER) of mature iPSC-RPE cells on transwells (4 replicates/genotype. 2-way ANOVA, ***p<0.001. Data represented as mean ± SD). **(E)** Quantification of FITC-labeled POS bound and **(F)** internalized by CRISPR-edited iPSC-derived RPE *PRPF31*^+*/*+^ and *PRPF31*^+*/-*^ cells at 30 min, 2 h, and 5 h (2-way ANOVA, ****p<0.0001, **p<0.01. Data represented as mean ± SD).

To test phagocytosis function of the *PRPF31*^+/-^ iPSC-RPE, CRISPR-edited *PRPF31*^+/+^ and *PRPF31*^+/-^ iPSC-RPE cells were incubated with POS for 30 min, 2 hours, and 5 hours. Analyses showed significant defects in both POS binding (2-way ANOVA, p=0.0001) and internalization (2-way ANOVA, p=0.0027) of POS in *PRPF31*^+/-^ cells compared to wild type at 2 hours (Figure 2E, F). Differences in binding and internalization were not significant at 30 minutes and 5 hours (Figure 2E, F).

### 2. Optimization of AAV vectors to be used for gene therapy in iPSC-RPE cells

Since *PRPF31*-associated retinal degeneration results from haploinsufficiency, AAV-mediated gene augmentation therapy can theoretically be used to rescue the retinal degeneration phenotype. Using mature iPSC-RPE at p2 on transwells, we tested a panel of AAV with a variety of serotypes (AAV2/2, AAV2/5, and AAV2/Anc80) expressing green fluorescent protein (EGFP) under the control of two different promoters: the regular cytomegalovirus (CMV), and the synthetic CASI promoter, which contains a portion of CMV, a portion of the chicken b-actin promoter, and a portion of the UBC enhancer ^28^. In addition, different concentrations of AAV were tested at 25,000, 50,000 or 100,000 viral genome copies per cell (GC/cell). To determine the highest transduction efficiency and stability, EGFP positive cells were counted at time points 72 hours, 2 weeks, 4 weeks, and 8 weeks post-transduction using fluorescence-activated cell sorting (FACS) (Figure 3). At 72 hours post transduction, cell toxicity was highest when transduced with the AAV2/5 serotype (2-way ANOVA. n=2 replicates/condition. p<0.0001). Toxicity for each AAV tested was increased at the higher titer, except for cells transduced with AAV2/Anc80, which did not show significant differences between 25,000 or 100,000 GC/cell (Figure 3A). AAV 2/2 and AAV2/Anc80 serotypes showed similar levels of long-term transduction, with the CASI promoter expressing higher levels of GFP versus CMV (2-way ANOVA. n=2 replicates/condition. p<0.0001) (Figure 3B). For the construct expressing the *PRPF31* gene, we chose the serotype AAV2/Anc80 with the CASI promoter, due to the lower cell toxicity at a higher titer, higher transduction expression of GFP, and high reported *in vivo* transduction efficiency ^29,30^. A detailed AAV-*PRPF31* vector map is provided in Figure S1.

**Figure 3.**
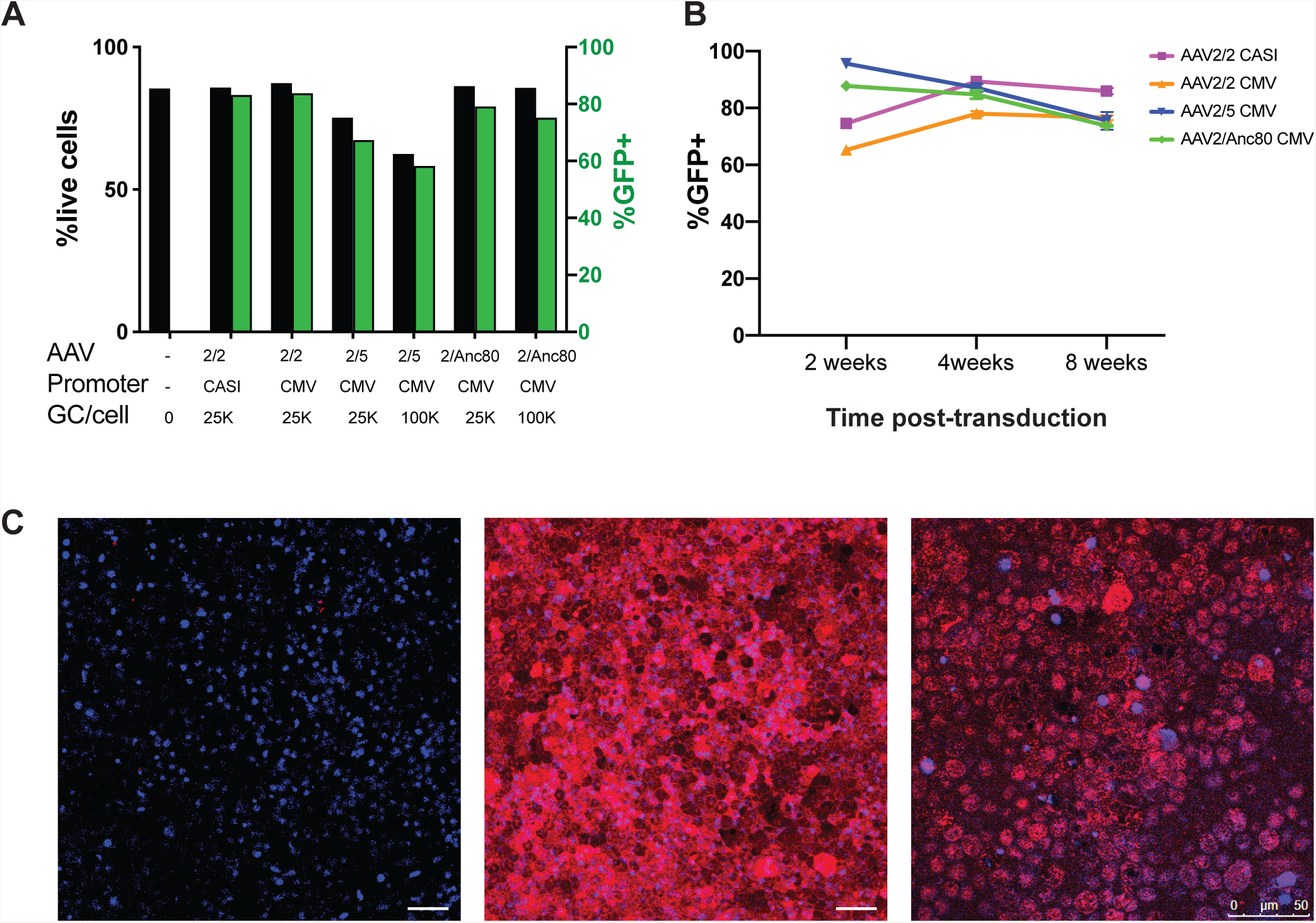
Optimization of AAV vectors. **(A)** Percentage of live cells (black bars) and GFP positive cells (green bars) 72 hours post transduction with AAV measured by FACS. **(B)** Percentage of GFP positive cells over the total population at 2, 4, and 8 weeks post-transduction with 25,000/cell (25k) AAV shows stable expression of GFP in iPSC-RPE cells over time. **(C)** RPE p2 transduced with 50,000 GC/cell (50k) AAV-*PRPF31* and immunostained with anti-V5 at 4 weeks post transduction. The third panel (higher magnification) shows the localization of the AAV-derived PRPF31 WT protein in the nuclei of the RPE cells. Scale bars= 50 µm.

### 3. Functional defects in mutant iPSC-RPE *PRPF31*+*/-*cells can be rescued with AAV-mediated gene augmentation

#### 3.1 Phagocytic defects in iPSC-RPE PRPF31+/-can be rescued by AAV-mediated gene augmentation

To test the use of gene augmentation therapy for *PRPF31*-associated disease, we treated patient derived and CRISPR-edited iPSC-RPE *PRPF31*^+/-^ cells and control cells with AAV-*PRPF31* ^12^. For these studies, differentiated iPSC-RPE cells at p2 were plated on matrigel-coated transwells and cultured in X-VIVO10 medium ^27^. After four weeks, cells were confluent and pigmented, and were transduced with several doses of *AAV2/Anc80 AAP.CASI.V5.PRPF31-mCHERRY.RBG* (from now, AAV-*PRPF31*). Cells were cultured for an additional four weeks after AAV-*PRPF31* treatment, and then *PRPF31* expression and phagocytosis activity were evaluated. Immunostaining with antibodies for the V5 tag demonstrated stable transduction after 4 weeks and correct localization of the AAV-derived PRPF31 protein in the nucleus (Figure 3). The phagocytic function of the AAV-*PRPF31* treated cells was assayed by challenging the cells with bovine FITC-labeled POS for 2 hours. Treatment with AAV-*PRPF31* resulted in rescue of binding and internalization efficiencies in iPSC-RPE cells *PRPF31*^+/-^ cells (2-way ANOVA, p=0.0322) (Figure 4).

**Figure 4.**
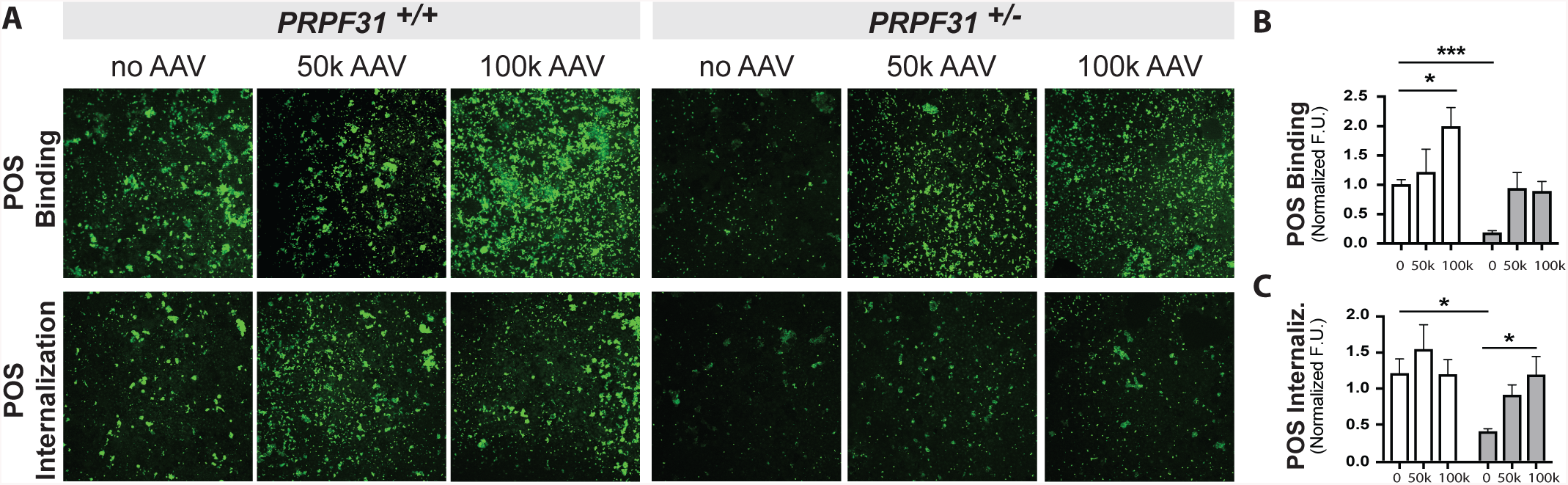
Phagocytosis defect is rescued by AAV-*PRPF31*. **(A)** Representative confocal fluorescent images (20x objective) and quantification of POS bound **(B)** and internalized **(C)** at 2h show significant phagocytosis defect in PRPF31 mutant cells *PRPF31*^+*/-*^ (white bars) compared to wildtype *PRPF31*^+*/*+^ (grey bars). Phagocytic function is restored in mutant cells after gene therapy with AAV-*PRPF31*. AAV 50K: 50,000 GC/cell, AAV 100K: 100,000 GC/cell. (n=2 replicates, 3fields/replicate, data represented as mean ± SEM. 2-way ANOVA, *p<0.05, ***p<0.001).

Interestingly, the phagocytic activity was enhanced in wild type iPSC-RPE *PRPF31*^+/+^ cells treated with AAV-*PRPF31* (Figure 4). Specifically, the capacity of binding POS increased by 2-fold when the cells were treated with 100,000 GC/cell (2-way ANOVA, p=0.0293) (Figure 4B), while the amount of internalized POS remained similar before and after AAV treatment (Figure 4C).

The effect of AAV-*PRPF31* treatment on phagocytosis function was also evaluated in patient derived iPSC-RPE *PRPF31*^+/-^ cells and corrected counterparts. The results of these studies demonstrated phagocytic activity was also restored in iPSC-RPE *PRPF31*^+/-^ cells from a different genetic background in a dose dependent manner (Figure 5).

**Figure 5.**
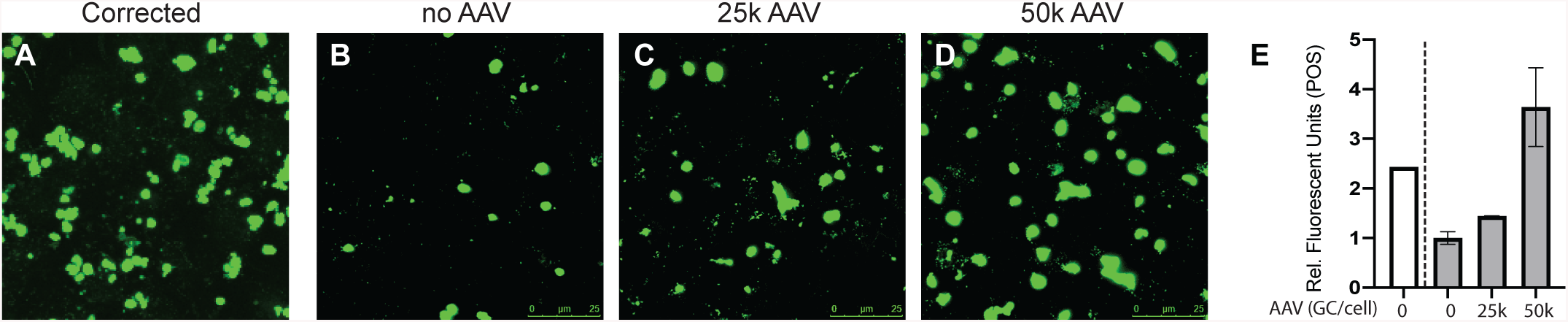
Patient-derived iPSC-RPE *PRPF31*^+*/-*^ cells show dose-dependent rescued phagocytic function after treatment with AAV-*PRPF31*. Confocal fluorescent images show FITC-labeled POS engulfed by the patient-derived iPSC-RPE p2 cells (corrected or mutant for PRPF31) on transwells treated with **(A, B)** no AAV, **(C)** 25,000 GC/cell AAV, or **(D)** 50,000 GC/cell AAV. Scale bars=25µm. **(E)** Fluorescence was quantified with ImageJ (n=2/treatment. One-way ANOVA, p=0.0655).

#### 3.2 Defects in cilia growth in iPSC-RPE PRPF31^+/-^ can be rescued by AAV-mediated gene augmentation

Among other defects, decreased cilia length was observed in patient derived *PRPF31*^+/-^ iPSC-RPE cells ^12^. Immunostaining with antibodies for the cilia-specific marker ARL13B showed similar results in CRISPR-edited iPSC-RPE *PRPF31*^+/-^ cells grown on transwells for 8 weeks, compared to wild type cells (Figure 6). To test the ability of gene augmentation to correct this cilia defect, patient derived and CRISPR-edited iPSC-RPE *PRPF31*^+/-^ cells were treated with AAV-*PRPF31*. Four weeks post-transduction with AAV-*PRPF31*, cilia length and incidence were restored to normal in both the patient-derived and CRISPR-edited iPSC-RPE *PRPF31*^+/-^ cells (Figure 6) (2-way ANOVA, n=50cells/culture. p<0.0001). No effect was observed in wild type cells treated with the AAV-*PRPF31* (Figure 6).

**Figure 6.**
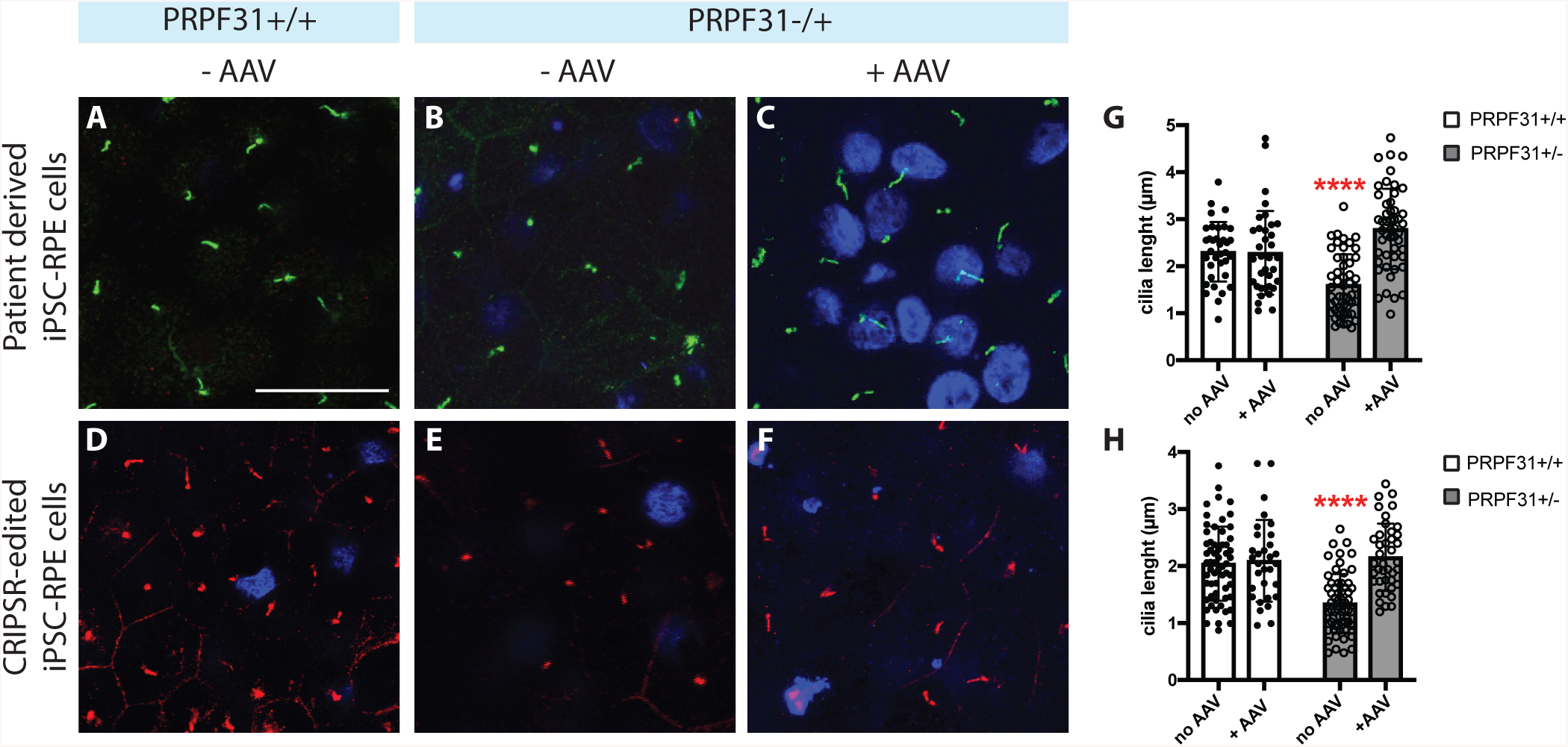
Shorter cilia length in iPSC-RPE *PRPF31*^+*/-*^ cells is rescued by AAV-*PRPF31*. **(A-F)** Immunostaining for ARL13B shows shorter cilia in both patient-derived and CRISPR edited iPSC-derived RPE *PRPF31*^+*/-*^ cells compared to the control. The length of the cilia is restored after transduction with 50k GC/cell of AAV-*PRPF31*. Due to the physical distance with the cilia, nuclei are out of plane in some of the images. Scale bar = 25µm. **(G-H)** Cilia length measured with Image J demonstrates significantly shorter cilia in *PRPF31*^+*/-*^ cells, which is rescued with AAV-based gene therapy. N=50 cells/culture. (Data represented as mean ± SD; individual cell measurement values are shown as boxes (wild type) or circles (*PRPF31*^+*/-*^) 2-way ANOVA, ****p<0.0001).

#### 3.3 Reduced barrier function in iPSC-RPE PRPF31^+/-^ cultures is partially rescued by AAV-PRPF31

The barrier function of the iPSC-RPE *PRPF31*^+/-^ cell monolayer was compromised, as indicated by lower levels of TER (Figure 2). To investigate the basis for this, we stained control and *PRPF31*^+/-^ mutant iPSC-RPE with antibodies to collagen VI (COL VI) to assess extracellular matrix elaboration. As shown in Figure 7, while the *PRPF31*^+/-^ cells remain polarized, with basolateral secretion of COL VI, they produce less of this extracellular matrix protein than control cells. Treatment of the iPSC-RPE *PRPF31*^+/-^ cells with AAV-*PRPF31* restored normal levels of COL VI production (Figure 7A-C).

**Figure 7.**
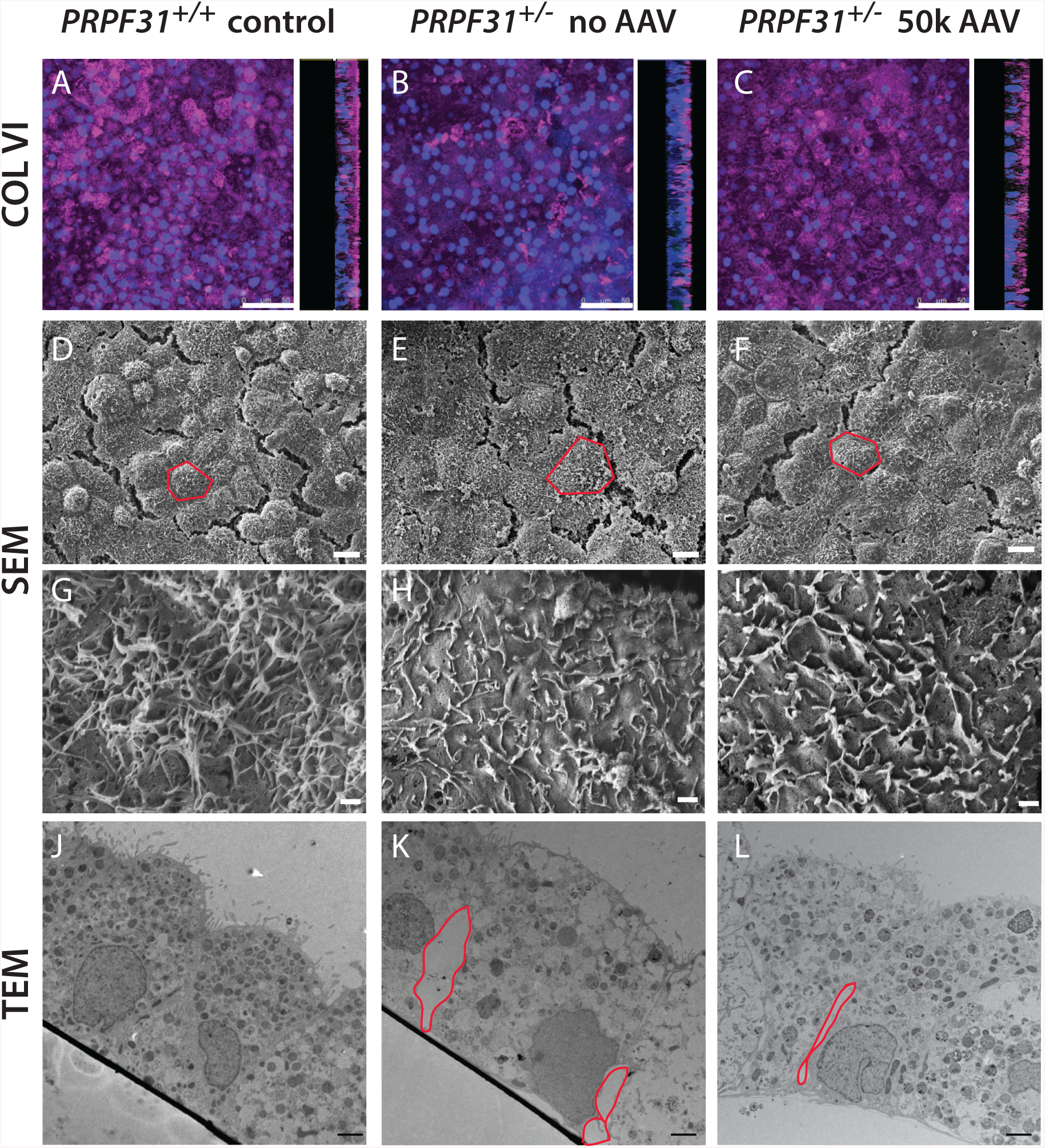
Confocal and electron micrographs of iPSC-derived RPE cell monolayer after 8 weeks on transwells. **(A)** Maximum intensity of COL VI expressed at 8 weeks. Z-stack was built from images taken every 0.13 μm with confocal microscope. 90° projections show that COL VI is deposited on the basal side of the RPE monolayer. Flat view by SEM at low **(D, E, F)** and high **(G, H, I)** magnification of *PRPF31*^+*/*+^ and *PRPF31*^+*/-*^ cell monolayers untreated or treated with AAV-*PRPF31* (single cells delimited by red lines). **(J, K, L)** Section view by TEM of *PRPF31*^+*/*+^ and *PRPF31*^+*/-*^ cell monolayers untreated or treated with AAV-*PRPF31* (red lines frame gaps between cells). Scale bars A-C: 50 µm, D-F: 10 µm, G-I: 1 µm, H-L: 2 µm.

Scanning electron microscopy (SEM) pictures revealed the presence of flatter and harder to distinguish cells in *PRPF31*^+/-^ cultures. The definition of the cells is partially restored following AAV treatment (Figure7 D-F). At higher magnification, *PRPF31*^+/-^ cells displayed fewer microvilli than controls with peculiarly coiled and flattened structure, which are also partially restored after treatment with AAV-*PRPF31* (Figure 7 D-I).

Disruption of the RPE monolayer in *PRPF31*^+/-^ cultures was confirmed by transmission electron microscopy (TEM), in which cross-section images showed gaps between adjacent cells at the basal side while apical tight-junctions stayed intact (Figure 7J, K), explaining the normal pattern of ZO-1 exhibited by iPSC-RPE mutant cells (Figure 2). Additionally, stress vacuoles and shorter microvilli were observed in the mutant iPSC-*PRPF31*^+/-^ cells. Treatment with AAV-*PRPF31* results in fewer stress vacuoles and reduced gaps between the RPE cells (Figure 7L). Despite improved RPE cell morphology, TER values were not restored in iPSC-*PRPF31*^+/-^ cultures after gene therapy treatment (Figure S2), possibly because basal gaps between cells, although smaller, were still present after treatment (Figure 7L).

## DISCUSSION

Autosomal dominant RP due to mutations in the *PRPF31* gene is caused by haploinsufficiency ^10,25^, providing an opportunity for treatment with gene-augmentation therapy. In this study, we modeled the cellular phenotype of the disease to test AAV-based *PRPF31* gene augmentation therapy. Mutant iPSC-RPE *PRPF31*^+*/-*^ cells grown on transwells for 8 weeks recapitulated the cellular phenotype associated with mutations in *PRPF31*, including abnormal structure, decreased phagocytic activity, defective ciliogenesis, and compromised barrier function. AAV-mediated *PRPF31* gene augmentation therapy restored normal phagocytosis function and cilia length and partially restored barrier function in patient-derived and genome-edited iPSC-RPE *PRPF31*^+*/-*^ cells. To the best of our knowledge, this is the first proof-of-concept that AAV-based gene therapy has the potential to be used for treatment of *PRPF31*-associated RP.

The RPE monolayer is the primary cell type affected in *PRPF31*-related RP, and a number of cell and animal models have been developed that recapitulate the *PRPF31*-associated cellular phenotypes ^6,9,12,25,31–33^. Consistent with prior reports, both the patient derived and genome edited *PRPF31* mutant iPSC-RPE cells used in the current study demonstrated defective phagocytosis and cilia formation ^12^. Our data suggest that mutations in *PRPF31* affect both binding and internalization of POS, with a larger impact in POS binding; analogous to the defect described in mutant mouse RPE *Prpf31*^+*/-*^ cells ^6^. Phagocytosis is a critical support function in continuously renewing light-sensitive outer segment portions, which is necessary for vision ^34^. With diminished phagocytic activity, individual POS components are not degraded or recycled back to photoreceptors, eventually leading to photoreceptor loss ^35^.

As expected by observations in RPE and optic cups derived from patients with mutations in *PRPF31* ^12^, iPSC-RPE *PRPF31*^+*/-*^ cells displayed shorter cilia. The cilium is critical for the RPE cell development and maturation, and for correct polarization of the RPE monolayer, which supports the photoreceptor integrity ^36^. Defects in the cilia of RPE *PRPF31*^+*/-*^ cells implies defects in maturation and polarization that compromise the function of the monolayer. A recent paper using ciliopathy patient derived iPSC-RPE cells established the maturation of the RPE cilium as the primary cause for disease ^36^. Interestingly, the authors demonstrated that treatment with modulators of ciliogenesis can rescue the phagocytosis activity by the RPE cells ^36^. Based on those results, we believe that the decreased phagocytosis activity observed in mutant iPSC-RPE *PRPF31*^+*/-*^ cells may be secondary to the abnormal maturation of cilia and microvilli. In any case, the degeneration of the RPE occurs first, later leading to degeneration of the neural retina observed in patients ^3,36^. Thus, we postulate that the RPE is the best candidate tissue to be targeted by gene therapy.

*In vivo*, RPE microvilli interdigitate with photoreceptors, providing mechanical support and executing the diurnal phagocytic removal of shed POS. Studies in mice have demonstrated that mutations in *Prpf31* resulted in diminished adhesion between RPE microvilli and POS, which delayed the phagocytic burst after light onset ^6^. Our cell-based model has allowed a better characterization of the microvilli structure in *PRPF31* mutant cells, showing deficiencies in microvilli length and frequency, as well as a coiled morphology, which may reduce POS binding to the integrin receptors, thus delaying phagocytic function. Additional structural defects displayed by the iPSC-RPE *PRPF31*^+*/-*^ cells were stress vacuoles and widened spaces between adjacent cells, which could be consequence of high apoptotic rates derived from aberrations in gene splicing. Splicing defects due to mutations in *PRPF31* have been specifically associated with functional and ultrastructural defects in the RPE ^12^. For instance, gaps between mutant cells could indicate reduced expression of focal adhesion genes; while diminished collagen may be associated with changes in regulators of extracellular matrix genes derived from mutations in *PRPF31* ^12^. Further transcriptome analyses are required to unravel the mechanisms underlying functional and structural defects in the RPE.

Patients with autosomal recessive IRD associated with mutations in RPE65 have been successfully treated and show improved visual function after sub-retinal injections with AAV expressing wild type RPE65 ^17^. Further, successful pre-clinical studies and clinical trials of gene therapies for other inherited retinal disorders are in progress ^18–21^. Likewise, we believe the visual function in patients with RP caused by *PRPF31* haploinsufficiency can be improved with AAV-mediated gene augmentation therapy. Based on this premise, we optimized an AAV vector that expresses wild type PRPF31 under the CASI promoter, which showed no toxicity in RPE cells. Although our experiments were performed *in vitro*, it is hoped the development of this therapy can be advanced so that it can be used clinically. To support this goal, the AAV vector used for these studies was generated with the Anc80 serotype, in which modifications in the viral capsid makes it more potent to transduce retinal tissue *in vivo,* as proven in mice and non-human primate studies ^29,30^.

The results presented here are a proof-of-concept that AAV-mediated PRPF31 expression can restore the phagocytic function and the cilia length of mutant RPE *PRPF31*^+*/-*^ cells. Additionally, AAV-*PRPF31* partially rescues the barrier function of the iPSC-RPE *PRPF31*^+*/-*^ cell monolayer, which displays tighter cell junctions and healthier RPE cells after treatment. Interestingly, AAV-mediated PRPF31 expression also enhances phagocytic activity in wild type RPE cells. Since AAV-based therapy is applied to pre-mature cells, this effect could imply an accelerated functional maturation of the RPE. Transcriptome analyses of these cells will contribute to understanding the mechanisms of RPE pathology in patients with *PRPF31*-linked RP.

It is also important to highlight that AAV-*PRPF31* transduced the RPE cells with high efficiency, which support its potential as a therapy for humans. However, based on the FACS analyses, there is a percentage of cells not targeted by the AAV, which will likely undergo apoptosis and prevent the TER values from being completely restored. These findings indicate that AAV-based gene therapy treatment may rescue the function of the RPE cells individually, while the global function of the RPE monolayer requires that most cells are targeted by the AAV. Further analyses are required to investigate whether earlier intervention with the AAV-*PRPF31* vector may result in a total rescue of the RPE barrier function. Still, the rescue provided by AAV-derived PRPF31 may be limited to certain RPE functions, while others like the barrier function cannot be rescued. Nonetheless, partial restoration of the RPE function and structure may be enough to preserve vision in patients with *PRPF31*-linked RP. Supplementary studies *in vivo* and using iPSC-derived optic cups may be warranted to identify the cellular targets for human therapy.

## MATERIALS AND METHODS

### Ethical Considerations

Patient derived samples used were obtained by Dr. Majlinda Lako’s lab, with informed consent according to the protocols approved by Yorkshire and the Humber Research Ethics Committee (REC ref. no. 03/362). Further information on the patients and controls is provided in Buskin et al. ^12^.

### Culture of iPSC

IMR90 hiPSCs were purchased from WiCell (Madison, WI, USA) and cultured on growth factor reduced (GFR) Matrigel basement membrane matrix (354230, Corning, Bedford, MA, USA) coated plates in mTeSR™1 (85850, STEMCELL Technologies, Cambridge, MA, USA). Patient-derived hiPSCs with a 1 bp deletion in PRPF31 and duplicate cells containing a CRISPR-Cas9 correction were generated in Dr. Lako’s lab (Institute of Genetic Medicine, Newcastle University) ^12,37–40^ and cultured similarly to IMR90 hiPSCs.

### Guide RNA design and CRISPR/Cas9 Genome Editing

The single guide RNA (sgRNA) target sequence (GCACCGCATCTACGAGTATG) was generated using the tool http://crispr.mit.edu/, with a score of 91. The sgRNA was cloned into the vector pSpCas9(BB)-2A-GFP (PX458) (a gift from Feng Zhang, Addgene plasmid no. 48138).

### Transfection of CRISPR-edited IMR90 iPSCs

Cells were transfected using the Amaxa Nucleofector Kit V (Lonza, Morristown, NJ, USA) per manufacturer instructions. Four micrograms of plasmid DNA were transfected per 10 million cells in a GFR-Matrigel coated 10 cm dish. Cells incubated undisturbed for 48 hours in mTeSR™ 1 with 1µM selective ROCK inhibitor (Y27632, Tocris, Minneapolis, MN, USA) to increase cell survival rate. After 48 hours, positively transfected cells were separated via fluorescence-activated cell sorting (FACS), due to transient GFP expression from the vector pSpCas9(BB)-2A-GFP (PX458). Positive clones were plated on GFR-Matrigel coated 10cm dishes in media composed of 50% fresh mTESR1 and 50% filtered conditioned media from confluent iPSC cultures until small colonies formed (roughly 8 days later). To prevent cross-contamination between clones, colonies were manually dissected and transferred to 1 colony/well in 96-well plates and expanded until 60-70% confluent (roughly 8 days later). Once split, three replica plates were created: one to keep cells growing in incubator, one to analyze genotype, and one to freeze clones ^41^.

Recent studies have raised the concern that Cas9 produces a p53-mediated DNA damage response in iPSCs, which reduces the editing efficiency ^42,43^. This finding suggests that defects in p53 improve the efficiency of genome editing in iPSCs, potentially leading to the generation of a biased cell population with defective p53 ^43^. To determine if this occurred during editing of *PRPF31*, PBS only or 1µg total of the p53 antagonist MDM2 (E3-204-050, R&D Systems, Minneapolis, MN, USA) in PBS was co-transfected with the plasmid DNA. No differences were observed in editing efficiency between cultures treated with or without rhMDM2 (Figure S3).

### Genotyping Edited Clones by Sanger Sequencing

DNA was extracted from the expanded cell clones using QuickExtract DNA extraction solution (QE0905T, Lucigen, Middleton, WI, USA). PCR fragments were amplified using primers F: 5’-GGACAAGTGCAAGAACAATGAGAACC-3’ and R: 5’-GGATGTAGCTTTCCCAAGGTCACAGTG-3’. Deletions were undetectable via gel electrophoresis, so PCR products were analyzed by Sanger Sequencing. Two mutant clones with 4 bp and 10 bp out-of-frame deletions and two wild-type clones with no detected abnormalities were expanded. Normal karyotypes of each iPSC line were validated with the hPSC Genetic Analysis Kit (07550, STEMCELL Technologies, Cambridge, MA, USA).

### Differentiation of iPSCs

Confluent cultures of iPSC were differentiated using the 14-day direct differentiation protocol previously described ^26,27^. Briefly, a given combination of Noggin, Dkk-1, Insulin Growth Factor 1, nicotinamide, Activin A, basic Fibroblast Growth Factor, SU-5402, and CHIR99021 produces directed differentiation of iPSC into RPE cells that exhibit key characteristics of the RPE, including pigmentation, honeycomb morphology, and expression of RPE markers. Every 30 days, cells were enzymatically digested with TryPLE Express (12604013, Thermo Fisher, Waltham, MA, USA), strained through a 40µm filter and seeded at a density of 10^5^ cells/cm^2^ onto Matrigel-coated plates in XVIVO-10 media. After 60 days in culture, cells were seeded onto Matrigel-coated 6.5mm polyestyrene transwells with 0.4µm pores (3470, Corning, Bedford, MA, USA) at a density of 20,0000 (low-density) or 200,000 (high-density) cells per Transwell. The CRISPR-edited PRPF31 mutant and wild-type clones with better overall RPE-like morphology were chosen for the remainder of the protocol.

### Quantitative Polymerase Chain Reaction (qPCR)

RNA from patient-derived iPSC and CRISPR-edited iPSC cells was extracted at passage 2 using the RNeasy Mini Kit (74104, Qiagen, Germantown, MD, USA). Next, cDNA was synthesized using random primers with the AffinityScript cDNA synthesis kit (600105, Agilent) and used to assay the expression of the RPE markers using the TaqMan probes described by Leach et al. ^27^ using the Stratagene Mx3005P qPCR system (Agilent, Santa Clara, CA, USA). Samples were run in triplicate using 10ng of cDNA per well and normalized to housekeeping genes using the delta-CT method. Statistical tests were performed with GraphPad Prism 7.

### Transduction of RPE cells

All RPE p2 cells matured for 4 weeks on transwells before transduction. *AAV2/Anc80 AAP.CASI.V5.PRPF31-mCHERRY.RBG* was suspended in X-VIVO (BE04-743Q, Lonza, Basel, Switzerland) with normocin (ant-nr-1, InvivoGen, San Diego, CA, USA) to achieve an MOI of 25,000, 50,000 and 100,000 vg/cell in final volume of 50µL for low density cultures or 100µl for high density cultures. AAV was added to the iPSC-RPE cells on transwells and incubated overnight. Media was changed next day and twice/week afterwards. All RPE cells were cultured for 4 weeks before performing phagocytosis assay.

### POS Phagocytosis Assay

Photoreceptor outer segments (POS) were purchased from InVision BioResources (98740, Seattle, WA, USA) and labeled with FITC for 1 hour at room temperature (F6434, Fisher, Waltham, MA, USA). POS were re-suspended in enough cell media to constitute 10 POS per cell. The patient-derived and CRISPR-edited iPSC-RPE cells were incubated with FITC-POS for 30 minutes, 2 hours, or 5 hours. At the end of the incubation period, FITC-POS were aspirated and samples washed with PBS thrice for 1 minute each to stop phagocytosis. To quench FITC-POS bound outside of the cell, half of the samples incubated with 0.4% Trypan blue in PBS for 10 minutes at room temperature. All samples challenged with FITC-POS were fixed with ice-cold 100% methanol for 10 minutes and rehydrated with PBS. Samples without FITC-POS were fixed with 4% Paraformaldehyde for 10 minutes, followed by 1% glutaraldehyde for 30 minutes, and left in PBS.

### Immunocytochemistry

Transwells were cut from the chamber, cut in half with a sharp blade, and cells permeabilized with 0.1% Triton X-100 for 15 minutes, then blocked with 0.1% BSA in PBS for 1 hour ^44^. Primary antibodies were diluted in 0.1% BSA in PBS and incubated with cells overnight at 4°. Primary antibodies used: ZO-1 (1:100, 61-7300, Invitrogen, Carlsbad, CA, USA), V5 (1:50, R96025, Invitrogen, Carlsbad, CA, USA), ARL13B (1:100, 17711-1-AP, Proteintech, Rosemont, IL, USA), and Collagen VI (1:100, AB6588, Abcam, Cambridge, MA, USA). Secondary antibody was diluted 1:500 in 0.1% BSA in PBS and incubated with cells for 2 hours at room temperature. Secondary antibodies used were labelled with: Alexa Fluor 488 (A11034, Thermo Fisher Scientific, Waltham, MA, USA), Alexa Fluor 555 (A21429, Thermo Fisher Scientific, Waltham, MA, USA), or Alexa Fluor 647 (A21235, Thermo Fisher Scientific, Waltham, MA, USA). Lastly, cells were incubated with DAPI, rinsed with PBS, mounted with Fluoromount G (17984-25, Electron Microscopy Sciences, Hatfield, PA, USA). Samples were imaged with TCS SP5 II confocal laser scanning microscope (Leica, Allendale, NJ, USA) or fluorescence microscope (Eclipse T, Nikon, Melville, NY, USA).

### Measurement of cilia length

Performed with Image J using scale bars to set the ratio pixel/micron.

### Quantification of the fluorescent signal

Images were converted to binary format with ImageJ. The integrated intensity was measured ^45,46^.

### Transmission Electron Microscopy (TEM)

Samples for TEM were prepared as previously described ^47^. Briefly, transwells were fixed with 2.5% glutaraldehyde overnight at 4° in 0.1M Cacodylate buffer. The transwell inserts containing cell monolayers were cut into smaller pieces, and post-fixed in 1.0% osmium tetroxide in cacodylate buffer for 1 hour at RT, then rinsed in cacodylate buffer. Insert pieces were then dehydrated through a graded series of ethanol, and placed pre-infiltrated overnight with propylene oxide and Eponate 1:1. Specimens were embedded in Eponate resin. 70 nm sections were cut using a Leica EM UC7 ultramicrotome, collected onto formvar-coated grids, stained with uranyl acetate and Reynold’s lead citrate and examined in a JEOL JEM 1011 transmission electron microscope at 80 kV.

### Scanning electron microscopy (SEM)

Transwell inserts containing exposed ECM after decellularization were fixed with 4% PFA for 10 min and 1% glutaraldehyde for 30 min at RT, followed by critical dehydration and Chromium coating as previously described ^46,47^. Samples were imaged by Field Emission Scanning Electron Microscope (JEOL 7401 F).

### Statistics

Results are expressed as mean ±SD, with p<0.05 considered statistically significant. Differences between groups were compared using the Student t-test or 2-way ANOVA as appropriate using the GraphPad Prism software.

## ACKNOWLEDGEMENTS

The authors would like to thank Dr. Michael Farkas and Maria Sousa for their preliminary studies, and Dr. Clegg and Leah Foltz for their technical support with the RPE differentiation method. We thank MEEI/SERI Gene Transfer Vector Core for viral vector production. We thank Diane Capen for her excellent technical assistance with the TEM, supported by the Microscopy Core of the Center for Systems Biology/Program in Membrane Biology, which is partially an Inflammatory Bowel Disease Grant DK043351 and a Boston Area Diabetes and Endocrinology Research Center (BADERC) Award DK057521. This work was supported by NIH Grant EY020902. The patient derived cells were generated with the support of RP Fighting Blindness (GR595), Fight for Sight (1456/1457), and ERC (CoG_614620). The authors declare no competing interest.

## AUTHOR CONTRIBUTIONS

Conceptualization, E.B., R.B., E.A.P, and R.F.G.; Methodology E.B, A.B., M.L., and R.F.G., Writing, Review and Editing, E.B., E.A.P, and R.F.G. Funding Acquisition, E.A.P. Supervision, E.A.P. and R.F.G.

