## Supplemental Figures for "AAV-mediated gene augmentation therapy restores critical functions in mutant iPSC-derived PRPF31^+/-^ cells"

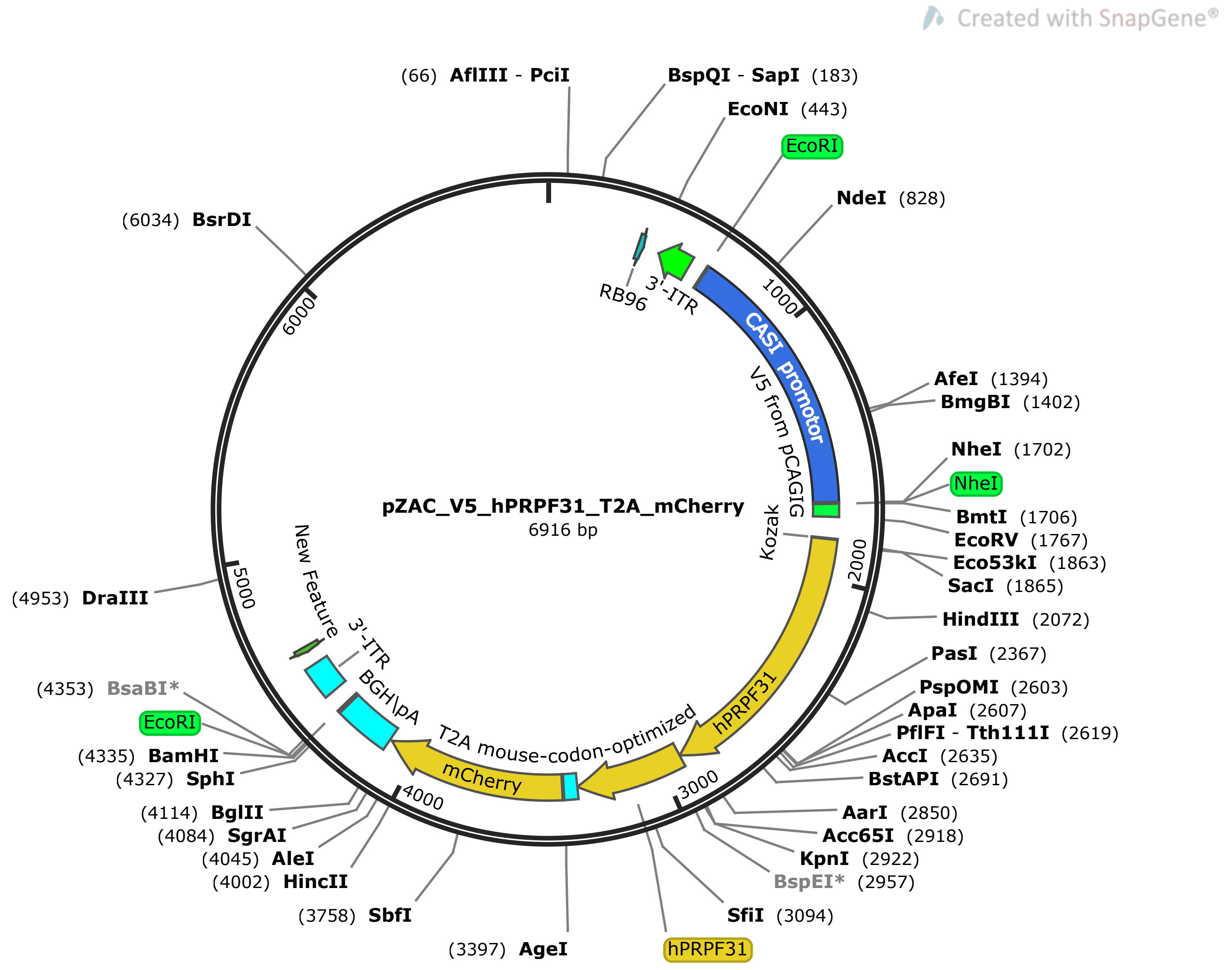


**Figure S1.** Map of the AAV vector used for AAV-mediated expression of PRPF31 in iPSC-RPE cells.

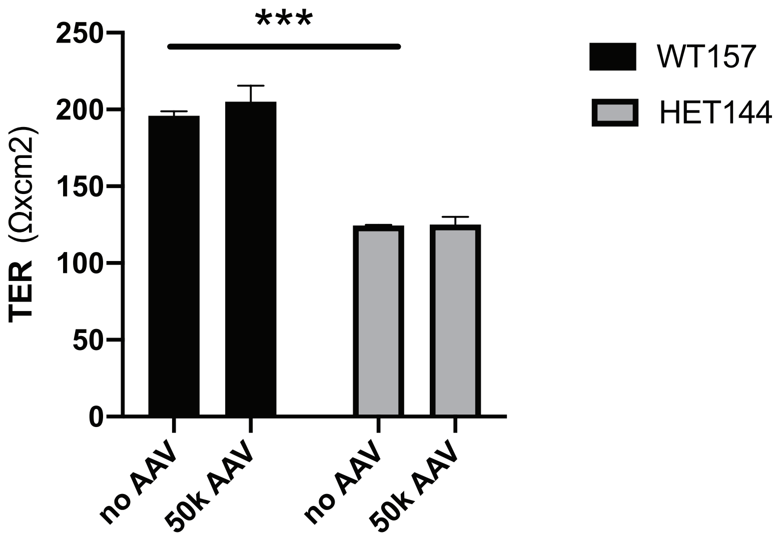


**Figure S2.** TER of iPSC-RPE PRPF31+/+ and PRPF31+/- cells on transwells with and without treatment with AAV-*PRPF31*. (n=4/type. 2-way ANOVA. ***p<0.001. Data represented as mean + SD).

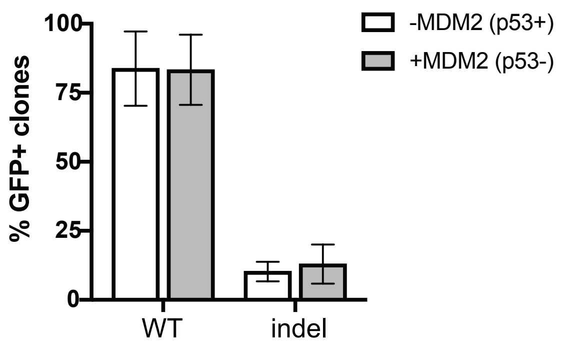


**Figure S3.** Percentage of GFP positive iPSCs with and without indels in PRPF31 after transfection with the plasmid containing Cas9 and the gRNA 1 in the absence (white bars) or presence (grey bars) of the p53 antagonist MDM2 (2-way ANOVA. Data represented as mean ± SD).
